# Variant analysis of SARS-CoV-2 genomes in the Middle East

**DOI:** 10.1101/2020.10.09.332692

**Authors:** Khalid Mubarak Bindayna, Shane Crinion

## Abstract

**Background:** Coronavirus (COVID-19) was introduced into society in late 2019 and has now reached over 26 million cases and 850,000 deaths. The Middle East has a death toll of ∼50,000 and over 20,000 of these are in Iran, which has over 350,000 confirmed cases. We expect that Iranian cases caused outbreaks in the neighbouring countries and that variant mapping and phylogenetic analysis can be used to prove this. We also aim to analyse the variants of severe acute respiratory syndrome coronavirus-2 (SARS-CoV-2) to characterise the common genome variants and provide useful data in the global effort to prevent further spread of COVID-19.

**Methods:** The approach uses bioinformatics approaches including multiple sequence alignment, variant calling and annotation and phylogenetic analysis to identify the genomic variants found in the region. The approach uses 122 samples from the 13 countries of the Middle East sourced from the Global Initiative on Sharing All Influenza Data (GISAID).

**Findings:** We identified 2200 distinct genome variants including 129 downstream gene variants, 298 frame shift variants, 789 missense variants, 1 start lost, 13 start gained, 1 stop lost, 249 synonymous variants and 720 upstream gene variants. The most common, high impact variants were 10818delTinsG, 2772delCinsC, 14159delCinsC and 2789delAinsA. Variant alignment and phylogenetic tree generation indicates that samples from Iran likely introduced COVID-19 to the rest of the Middle East.

**Interpretation:** The phylogenetic and variant analysis provides unique insight into mutation types in genomes. Initial introduction of COVID-19 was most likely due to Iranian transmission. Some countries show evidence of novel mutations and unique strains. Increased time in small populations is likely to contribute to more unique genomes. This study provides more in depth analysis of the variants affecting in the region than any other study.

**Funding:** None

## Introduction

On January 9^h^ 2020, the China Centre for Disease Control reported that 15 of 59 suspected cases of pneumonia were due to a novel human coronavirus (CoV), now known as Severe Acute Respiratory Syndrome CoV 2 (SARS-CoV-2) ^1^. The genome for this novel virus was then made publicly available on the *Global Initiative on Sharing All Influenza Data (GISAID)* the next day. SARS-CoV-2 is an easily spreadable virus which would evolve into a global pandemic of at least 26 million cases and 850,000 deaths^2^. One of the first countries to experience a significant outbreak was Iran. The country reported its first confirmed case on 19^th^ February 2020 from a merchant in Qom who travelled from China^3.^. Many of the first countries with infections in the Middle East were linked to travellers from Iran including Lebanon, Kuwait, Bahrain, Iraq, Oman and UAE. COVID-19 continued to spread to the remaining Middle Eastern countries with a death toll of over 50,000 people according to health authorities. This number is expected to be an underestimation due to countries effected by war including Libya, Syria and Yemen. Needless to say, there have been devastating effects to the region and the real effects are expected to be unreported^4^.

Researchers are racing to develop a vaccine that can provide viral immunity and avoid additional deaths. SAR-CoV-2 is transmitted using the spike protein which binds to human angiotensin-converting enzyme 2 (ACE2) receptor; the virus is easily transmittable due to mutations in the receptor-binding (S1) and fusion (S2) domain of the strain^5^. Transmission could be made even easier if more mutations accumulate. Although mutations are rare, they can create new strains and it is not guaranteed that the current leading vaccine trials will be effective as SARS-CoV-2 continues to mutate^6^. By categorizing variants, we can identify any new strains and how the mutations are likely to affect spread. As the Middle East is often under reported, it is important to characterise the variants of strains that are commonly present. Analysis of the common variants in the Middle East is essential to develop a vaccine that treats the strains in the region. This analysis helps understand the viral genome landscape and identify clades of the region.

## 2. Objectives

- Our hypothesis is that variants found in SARS-CoV-2 genomes from Middle Eastern samples will indicate delivery from Iran. We will use bioinformatics tools and publicly available samples to explore the composition of strains within each country. We expect that many strains will show evidence of Iranian origin.
- The aim is to explore the structure of Middle Eastern genome strains using multiple sequence alignment, tree generation and variant prediction (and others). If we explore the structure and common variants of SARS-CoV-2 strains in these populations, we expect to learn more about how the virus spread.

## 3. Methods

### Sample Source

We obtained the publicly available data from the Global Initiative on Sharing All Influenza Data (GISAID)^7^.

### Sample Size

122 Middle Eastern samples, Wuhan reference sequence NC_045512 and 5 recent Wuhan samples.

### Sample Selection

Samples were selected from the Middle East by using filtering available on the GISAID website. Only complete genome samples were used. The countries considered were Afghanistan, Bahrain, Cyprus, Egypt, Iraq, Iran, Israel, Jordan, Kuwait, Lebanon, Libya, Oman, Qatar, Saudi Arabia, Sudan, Syria, Turkey, United Arab Emirates (UAE) and Yemen. Iran had only 7 samples available after filtering for low coverage. Cyprus, Kuwait, Lebanon all had 8 samples available after filtering. No samples were available on the database from Afghanistan, Iraq, Libya, Sudan, Syria or Yemen. 10 samples were taken from all other countries. Samples were also filtered to high quality when possible. 10 samples was selected as the optimum number to cover all possible countries and remain within alignment file limit of size 4 Mb (maximum size for Clustal Omega tool). In countries with 10 samples, to prevent sample sourcing from same outbreak, the 5 earliest and 5 most recent samples were taken. All samples were downloaded from GISAID and then concatenated into a single multi-sample file and saved in FASTA format.

### Multiple sequence alignment

Using the collected samples, multiple sequence alignment (MSA) was performed using Clustal Omega (https://www.ebi.ac.uk/Tools/msa/clustalo/)^8^. The Clustal Omega online tool was used to perform the alignment (found at: https://www.ebi.ac.uk/Tools/msa/clustalo/). The online tools allows up to 4000 sequences or a maximum file size of 4 MB, therefore the maximum number of samples was used. The concatenated file of samples was uploaded to the online tool. For step 2, the output parameters selected were PEARSON/FASTA. All other options were kept at the default option. The output file generated is an alignment file; the file consists of all sequences with gaps denoted by ‘-’. The output file format is also a FASTA file.

### Variant identification

Variant calling was performed using the alignment FASTA file and the SNP extraction tool snp-sites^9^ (https://github.com/sanger-pathogens/snp-sites). These tools identify the SNP sites by taking a multi-sample FASTA file as input. The program then restructures the data as a variant call format (VCF) file. The VCF file provides a clear mapping of SNPs from the aligned sequences – this allows us to easily identify the SNP location and the genotype for each sample at a given locus. In the outputted VCF file, the rows correspond with each unique variant and the column provides the genotype at the given site.

### Variant annotation

We used SNP-eff^10^ to perform the variant annotation information such as the variant definition and the overlapping gene (found at: (https://pcingola.github.io/SnpEff/SnpEff.html). SnpEff also predicts the effect of the variants. SNPeff is integrated into the Galaxy web-based tool for bioinformatics analysis (found at: usegalaxy.org) ^11^. We utilised the Galaxy platform and uploaded our VCF file to Galaxy using their online upload tool. To annotate variants, you must first build a database from the reference genome. This is performed using the “SnpEff build” tool on Galaxy. To create the SnpEff database, we downloaded data from NCBI for Wuhan reference NC_045512.2 and uploaded to Galaxy. To build the database, we directed to NCBI and searched for NC_045512. We then downloaded the corresponding GFF file, which contains the annotations, and the FASTA file, which contains the entire genome. We then selected the build database option. Once the database was built, we selected the “SnpEff eff” tool to annotate variants. Galaxy populates the fields for VCF with the uploaded file from the previous step. The output format is selected as VCF and CSV report was also selected for additional useful information for downstream analysis. For Genome source parameter, the option “Custom” was selected to use the newly created database. Other default parameters were used including 5000 bases for Upstream / Downstream length and 2 bases as set size for splice sites (donors and acceptor) in bases. All filter output and additional annotation options were deselected and analysis was ran. Once the analysis was executed, the annotation data is outputted as an annotated VCF and a HTML report file.

### Data visualisation

Once the annotated VCF was generated, the VCF was imported to R for extraction of the variant annotation information. The annotated data was imported, manipulated and plotted using R v3.6.2^12^. The dplyr v0.8.4 package was used to summarise and align the data^13^. The ggplot2 package was used to align the identified variants and visualise the types of mutations that re-occur^14^. The x-axis in the plots indicates the variant position along the SARS-CoV-2 genome; the left y-axis indicates the sample name and the right y-axis represents the country of origin for each sample. This plot is used to compare the genome in different populations. The data was then ordered by date of first reported case, meaning that Wuhan is followed by United Arab Emirates and the final country is Cyprus.

### Phylogenetic analysis

Phylogenetic analysis was performed using BEAST (Bayesian Evolutionary Analysis Sample Trees) v1.10.4, to perform Bayesian analysis of molecular sequences using Monte Carlo Markov Chains (MCMC)^15^. The analysis followed the approach recommended to reconstruct the evolutionary dynamics of an epidemic. The aim of this is to obtain an estimate of the origin of the epidemic in the region and understand how it spread through the Middle East. To undertake the analysis, we opened BEAUTi, the graphical application used to analyse the control file. Although it requests a NEXUS file, the FASTA file can also be used. The data was uploaded using the Import Data option and appeared under the Partitions section. BEAUTi confirmed that 30851 sites are present in the uploaded data. The default options are selected for site model and clock model. Next, we specified the individual virus dates by selecting the “Tips” panel and selecting the “Use tip dates” option. A tab delimited file was uploaded which specified the upload date. This information was extracted from the names as they were downloaded from GISAID. Next, we set the substitution model by selecting the “Sites” tab and selected the default options of HKY model, the default Estimated base frequencies and select Gamma as Site Heterogeneity Model. Next, the molecular clock was selected under the “Clock” tab as a strict clock since we know that the frequency of mutation is low. The tree options are elected under the “Tree Prior” tab as “Random starting tree” for the tree model and “Coalescent: Exponential Growth”, a model that assumes a finite but constant population size and predicts that all alleles will be removed from the population individually. This provides additional predictions on the reproductive rate. In the “Priors” tab, select the scale as 100 for prior distribution which models the expected growth for a pandemic. The operators require no changes from the default. The MCMC option for chain length is set as 100,000 and sampling frequency to 100.

### Tree visualisation

Finally, we summarised the tree using the TreeAnnotator tool, an additional package as part of BEAST. We first select the file generated using BEAST and outputted the tree file. Then the output NEXUS file was imported to FigTree program to display. Once we opened FigTree to display, we re-ordered the order by increasing value and then switched on Branch Labels. We switched on Node Bars and selected the 95% highest posterior density (HPD) credible intervals for the node heights. We plotted a time scale by turning on the Scale Axis and then setting the Time Scale section for Offset as 2020.7, the latest date of collection for our samples.

## 4. Results

Sequence alignment and variant calling were completed successfully. Once these were complete, variation annotation was performed. We identified 2200 distinct genome variants which are recorded in Table 1. The most common, high impact variants were 10818delTinsG, 2772delCinsC, 14159delCinsC and 2789delAinsA. The frequency of each unique variant type can be found in Table 2, which outlines the locus of all SNPs with over 50 instances.

**Table 1:**
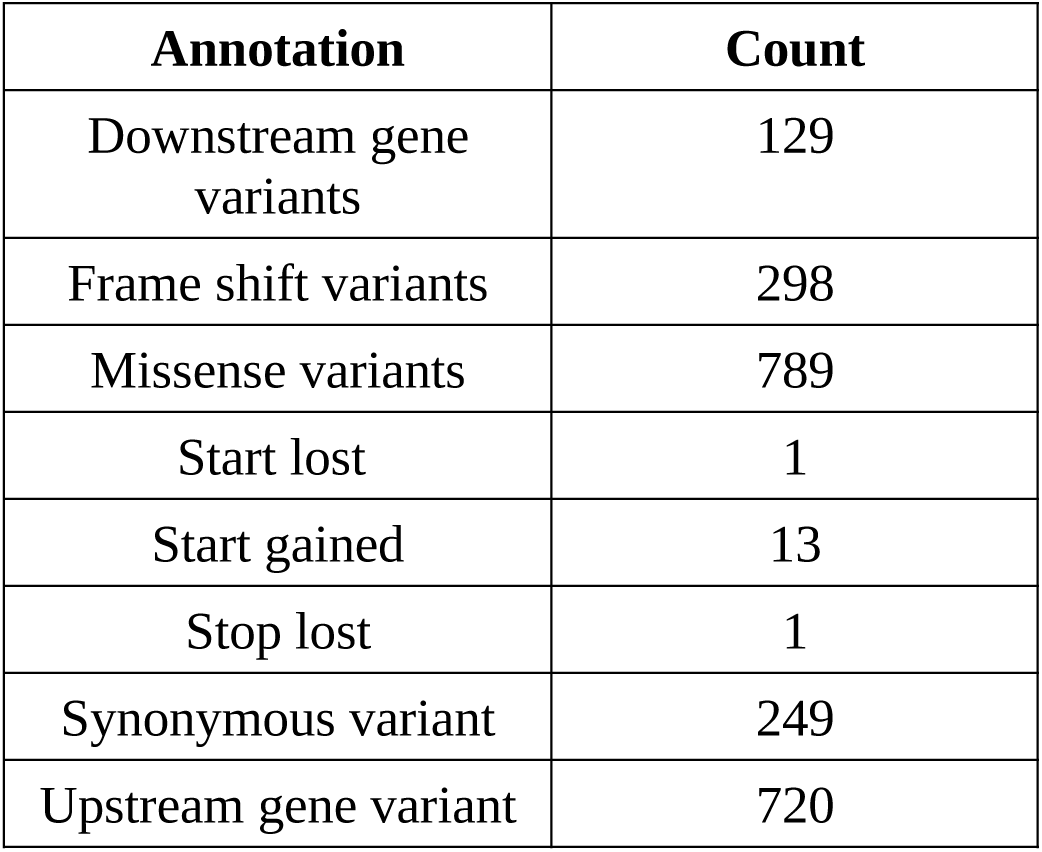
The frequency of each type of mutation found in the data.

**Table 2:**
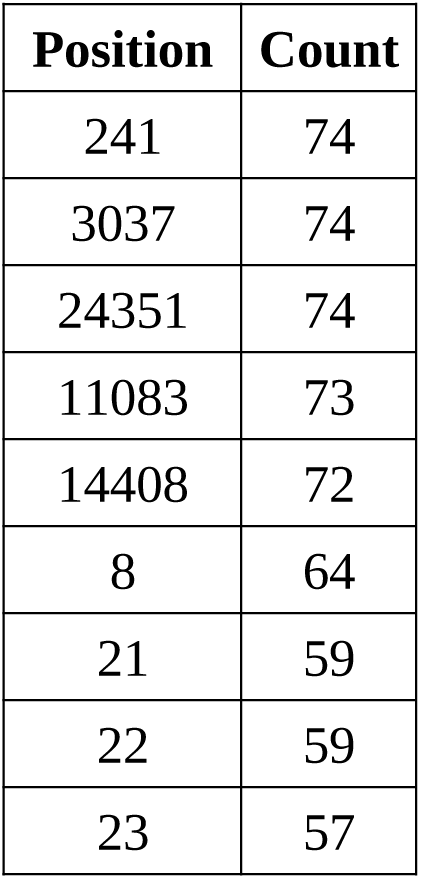
The frequency of mutations at each locus with > 50 hits.

**Table 3:** Catalogue of sample accession ID by country.

|  | **Region** | **ID** |
| --- | --- | --- |
| 1 | Bahrain | EPI_ISL_487274 |
| 2 | Bahrain | EPI_ISL_486889 |
| 3 | Bahrain | EPI_ISL_483545 |
| 4 | Bahrain | EPI_ISL_510528 |
| 5 | Bahrain | EPI_ISL_483542 |
| 6 | Bahrain | EPI_ISL_483548 |
| 7 | Bahrain | EPI_ISL_483547 |
| 8 | Bahrain | EPI_ISL_483543 |
| 9 | Bahrain | EPI_ISL_485401 |
| 10 | Bahrain | EPI_ISL_487273 |
| 11 | Cyprus | EPI_ISL_463742 |
| 12 | Cyprus | EPI_ISL_463743 |
| 13 | Cyprus | EPI_ISL_463744 |
| 14 | Cyprus | EPI_ISL_463748 |
| 15 | Cyprus | EPI_ISL_463741 |
| 16 | Cyprus | EPI_ISL_463745 |
| 17 | Cyprus | EPI_ISL_463746 |
| 18 | Cyprus | EPI_ISL_463747 |
| 19 | Egypt | EPI_ISL_482761 |
| 20 | Egypt | EPI_ISL_479735 |
| 21 | Egypt | EPI_ISL_479733 |
| 22 | Egypt | EPI_ISL_479734 |
| 23 | Egypt | EPI_ISL_510532 |
| 24 | Egypt | EPI_ISL_430819 |
| 25 | Egypt | EPI_ISL_430820 |
| 26 | Egypt | EPI_ISL_479732 |
| 27 | Egypt | EPI_ISL_482759 |
| 28 | Egypt | EPI_ISL_482760 |
| 29 | Iran | EPI_ISL_424349 |
| 30 | Iran | EPI_ISL_445088 |
| 31 | Iran | EPI_ISL_507007 |
| 32 | Iran | EPI_ISL_514753 |
| 33 | Iran | EPI_ISL_437512 |
| 34 | Iran | EPI_ISL_442044 |
| 35 | Iran | EPI_ISL_442523 |
| 36 | Israel | EPI_ISL_435291 |
| 37 | Israel | EPI_ISL_514303 |
| 38 | Israel | EPI_ISL_514305 |
| 39 | Israel | EPI_ISL_514302 |
| 40 | Israel | EPI_ISL_435286 |
| 41 | Israel | EPI_ISL_435289 |
| 42 | Israel | EPI_ISL_419211 |
| 43 | Israel | EPI_ISL_435284 |
| 44 | Israel | EPI_ISL_514306 |
| 45 | Israel | EPI_ISL_514301 |
| 46 | Jordan | EPI_ISL_430012 |
| 47 | Jordan | EPI_ISL_430002 |
| 48 | Jordan | EPI_ISL_430003 |
| 49 | Jordan | EPI_ISL_430009 |
| 50 | Jordan | EPI_ISL_429993 |
| 51 | Jordan | EPI_ISL_450188 |
| 52 | Jordan | EPI_ISL_434516 |
| 53 | Jordan | EPI_ISL_429997 |
| 54 | Jordan | EPI_ISL_450186 |
| 55 | Jordan | EPI_ISL_450187 |
| 56 | Kuwait | EPI_ISL_421652 |
| 57 | Kuwait | EPI_ISL_422427 |
| 58 | Kuwait | EPI_ISL_416543 |
| 59 | Kuwait | EPI_ISL_416458 |
| 60 | Kuwait | EPI_ISL_416541 |
| 61 | Kuwait | EPI_ISL_416542 |
| 62 | Kuwait | EPI_ISL_422426 |
| 63 | Kuwait | EPI_ISL_422424 |
| 64 | Lebanon | EPI_ISL_498551 |
| 65 | Lebanon | EPI_ISL_498552 |
| 66 | Lebanon | EPI_ISL_498554 |
| 67 | Lebanon | EPI_ISL_450512 |
| 68 | Lebanon | EPI_ISL_450515 |
| 69 | Lebanon | EPI_ISL_450511 |
| 70 | Lebanon | EPI_ISL_450508 |
| 71 | Lebanon | EPI_ISL_450509 |
| 72 | Oman | EPI_ISL_492023 |
| 73 | Oman | EPI_ISL_492026 |
| 74 | Oman | EPI_ISL_492024 |
| 75 | Oman | EPI_ISL_492065 |
| 76 | Oman | EPI_ISL_492025 |
| 77 | Oman | EPI_ISL_457706 |
| 78 | Oman | EPI_ISL_457704 |
| 79 | Oman | EPI_ISL_457937 |
| 80 | Oman | EPI_ISL_457701 |
| 81 | Oman | EPI_ISL_457974 |
| 82 | Qatar | EPI_ISL_427404 |
| 83 | Qatar | EPI_ISL_427420 |
| 84 | Qatar | EPI_ISL_427407 |
| 85 | Qatar | EPI_ISL_427406 |
| 86 | Qatar | EPI_ISL_427419 |
| 87 | Qatar | EPI_ISL_427418 |
| 88 | Qatar | EPI_ISL_427405 |
| 89 | Qatar | EPI_ISL_427408 |
| 90 | Qatar | EPI_ISL_427417 |
| 91 | Qatar | EPI_ISL_427416 |
| 92 | SaudiArabia | EPI_ISL_512924 |
| 93 | SaudiArabia | EPI_ISL_512926 |
| 94 | SaudiArabia | EPI_ISL_489996 |
| 95 | SaudiArabia | EPI_ISL_489998 |
| 96 | SaudiArabia | EPI_ISL_490000 |
| 97 | SaudiArabia | EPI_ISL_489999 |
| 98 | SaudiArabia | EPI_ISL_489997 |
| 99 | SaudiArabia | EPI_ISL_512927 |
| 100 | SaudiArabia | EPI_ISL_512922 |
| 101 | SaudiArabia | EPI_ISL_512923 |
| 102 | Turkey | EPI_ISL_495421 |
| 103 | Turkey | EPI_ISL_495445 |
| 104 | Turkey | EPI_ISL_495433 |
| 105 | Turkey | EPI_ISL_429868 |
| 106 | Turkey | EPI_ISL_429867 |
| 107 | Turkey | EPI_ISL_429866 |
| 108 | Turkey | EPI_ISL_428712 |
| 109 | Turkey | EPI_ISL_495436 |
| 110 | Turkey | EPI_ISL_495429 |
| 111 | Turkey | EPI_ISL_424366 |
| 112 | UnitedArabEmirates | EPI_ISL_469277 |
| 113 | UnitedArabEmirates | EPI_ISL_469279 |
| 114 | UnitedArabEmirates | EPI_ISL_435126 |
| 115 | UnitedArabEmirates | EPI_ISL_435121 |
| 116 | UnitedArabEmirates | EPI_ISL_435131 |
| 117 | UnitedArabEmirates | EPI_ISL_463740 |
| 118 | UnitedArabEmirates | EPI_ISL_469281 |
| 119 | UnitedArabEmirates | EPI_ISL_469280 |
| 120 | UnitedArabEmirates | EPI_ISL_435137 |
| 121 | UnitedArabEmirates | EPI_ISL_435134 |
| 122 | Wuhan | EPI_ISL_402123 |
| 123 | Wuhan | EPI_ISL_454949 |
| 124 | Wuhan | EPI_ISL_454948 |
| 125 | Wuhan | EPI_ISL_406798 |
| 126 | Wuhan | EPI_ISL_454951 |
| 127 | Wuhan | EPI_ISL_403931 |

These results were then used to generate the dotplot of variant alignment (Figure 1). The dotplot successfully indicated a pattern in variants that could not be easily identified from the alignment or annotation files. The alignment includes samples in facets based on their country. The alignment shows a pattern in variants that occur between each country. For example, this is prominent in Qatar, Jordan and Oman where the pattern makes the country distinctive from the variants plotted for other countries. In addition, the phylogenetic tree generated branching indicative of an Iranian origin (Figure 2)

**Figure 1:**
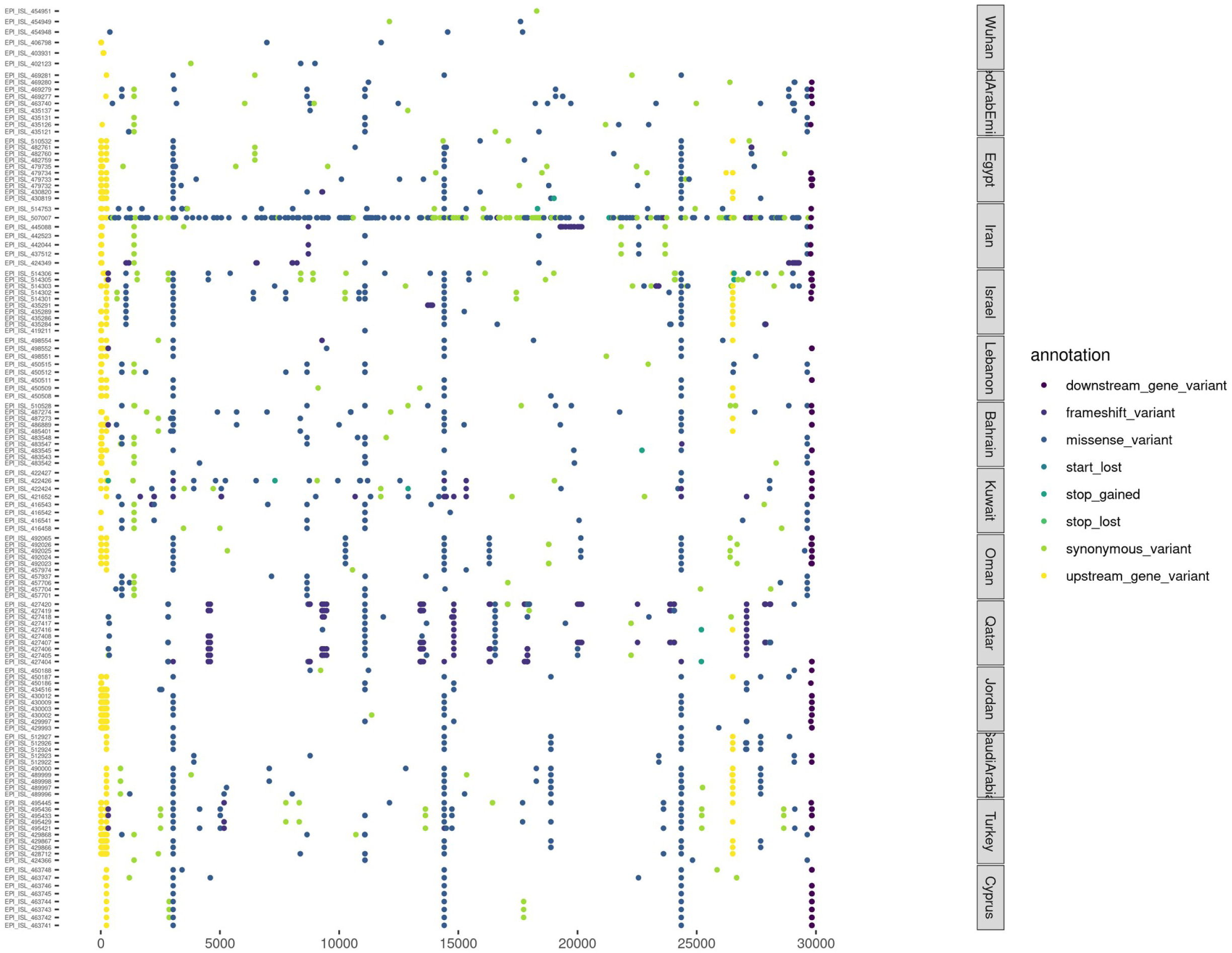
Dotplot of variants per sample by country. All samples included in the above plot are grouped by country of origin. The left x-axis denotes the accession ID and the right x-axis denotes the country of origin. The y-axis is the position along the SARS-CoV-2 genome. The order of countries is based on date of first reported of COVID-19. All variants are in relation to Wuhan reference sequence NC_045512.2. The Wuhan samples are in the top facet and show low mutation frequency in comparison to the reference. The following samples show a greater accumulation of mutations. One Iranian sample has many more variants than orders. The Saudi Arabia also appears to have a low mutation rate which may be due to their early contraction of the virus. Oman samples also indicate evidence of a distinctive strain. Many cases feature a missense variant at 3037, 14408 and 24351 base pairs.

**Figure 2:**
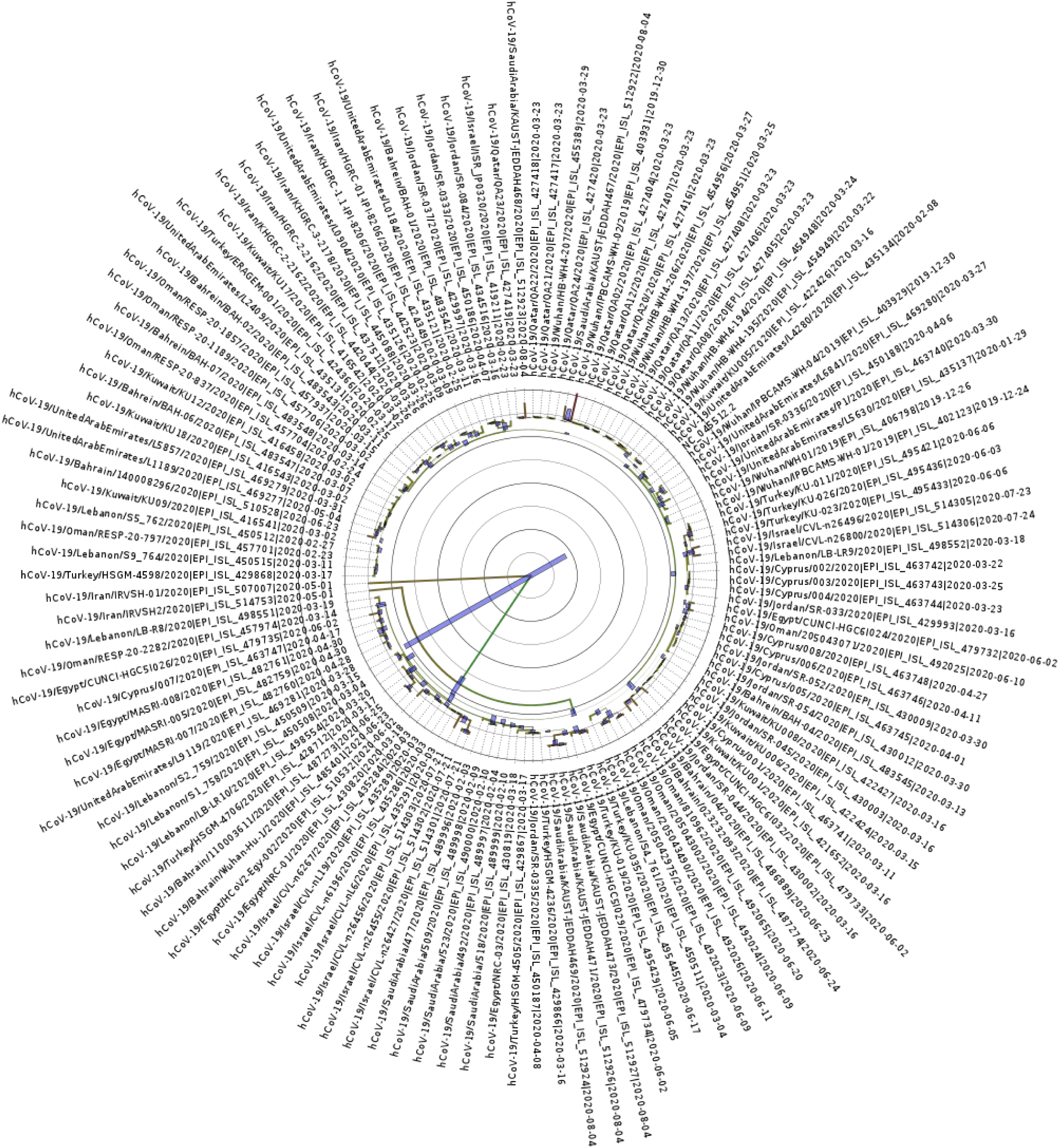
Phylogenetic tree for the SARS-CoV-2 genomes. Contains all 128 samples (including reference NC_045512). The colours represent the greatest height in association. The blue lines represent the 95 % HDP for each region. The highest branching point is with Iran samples EPI_ISL_507007 and EPI_ISL_514753 with length of 4.319 and 0.8752 respectively.

## 5. Discussion

The aim of this study was to identify whether COVID-19 was introduced to the Middle East from Iran and also to explore the genomic composition in the region.

Our study performs sequence alignment to compare all sequences against the reference genome. Once this is complete, the annotated variants were extracted to generate a plot mapping variants, grouping samples by country. The plot as seen in Figure 1, shows clear distinctive patterns within countries that are not obvious from the generated alignment and annotation files. The variants found in different regions, ordered by their first reported case, are from United Arab Emirates (UAE), Egypt, Iran, Israel, Lebanon, Bahrain, Kuwait, Oman, Qatar, Jordan, Saudi Arabia, Turkey and Cyprus. The variants at position 241, 3037, 24351, 11083 appear in many Middle Eastern countries but do not occur in Wuhan samples. This variant characterisation may be useful in the fight against COVID-19 and the development of treatments. Identifying unique variants to a region may explain why treatment is working for some and not others, should the mutations have an effect on the delivery or the severity of the virus.

The Iranian samples appear more diverse and interestingly do not share the mutation at 14408 which is evident in most samples. Iranian sample EPI_ISL_507007 appears to have a high frequency of variants that is not seen in others. THE GISAID detection system indicates no faults were found with sample EPI_ISL_507007 and report a full sequence match so it was not removed. It is noteworthy that the Iran samples appear to have a lower average SNP frequency than other countries. This may indicate that the virus transmitted to Iran earlier than other countries, as we expected. Israel, Bahrain, Kuwait, Oman, Saudi Arabia, Jordan and Turkey all share similar variant mapping. Qatar shows an usual mapping with a high frequency of frameshift variants. This may indicate a diverse, new strain is circulating in the country. Cyprus has little diversity in the variant mapping which is surprising given its late date for first reported cases. Another interesting point is that time-varied samples were taken for countries with 10 samples. We see not indication that there are distinct groups within countries. This further indicates the the mutation frequency is low. It also indicates that there is more variation in the genomic composition in samples from different countries than differences found in samples from different collection times. Smaller populations can cause greater accumulation of variants through genetic drift. This may occur given local lockdowns and travel restrictions that have been enforced worldwide. It is possible that these genomic strains with new mutations may create a situation where the countries develop a deathly strain that is not prominent in other parts of the world. This could result in a situation where a country is disproportionately affected by accumulating deaths or an inefficient vaccine.

Phylogenetic trees help in understanding the evolutionary relationships between groups. In the present context, we use them to identify the earliest strains and to track the spread of COVID-19 across the Middle East. The tree shows that UAE samples are distinguished and form one clade. This correlates with their early intervention and lockdown and subsequently appears to have resulted in a unique genome.

Samples from Qatar also form the majority of 1 clade, with many of the Wuhan samples, indicating that they are similar to the Wuhan samples and show little distinction. Egypt also becomes a distinct branch earlier than most samples. These examples are indicative of the global response – the lockdown of each country and prevention of spreading has resulted in SARS-CoV-2 strains of great similarity within each country. If lockdowns were not enforced, it is likely that these clades would be less distinguisable as mutations are spread between countries. As we expected, 2 of the highest branches points attach to Iranian samples, further implementing Iran in the initial spread across the Middle East. The phylogenetic tree therefore indicates what we suggest in our hypothesis – most samples originate from the Iranian sample. This is not surprising given the vast number of cases and early crisis state of the country. However, it is useful to see that the variant analysis shows what we suspect at the genome level. A related study also came to this conclusion by using contact tracing from cases related to religious events in the city of Qom, Iran^16^.

## Acknowledgements

The authors are grateful for the timely sequencing and release of genomes to make this study possible and to Dr. Anusha C P for her comments.

